# A comparative study of the chemotoxic effects of idarubicin and doxorubicin in a chemically-induced *in vivo* mouse model and in vitro models for hepatocellular carcinoma

**DOI:** 10.1101/2025.03.14.643356

**Authors:** Ada Lerma-Clavero, Maria Kopsida, Nathalie Arendt, Hans Lennernäs, Markus Sjöblom, Femke Heindryckx

**Affiliations:** Department of Medical Cell Biology, Uppsala University, Uppsala, Sweden; Department of Pharmaceutical Biosciences, Uppsala University, Uppsala, Sweden

## Abstract

Hepatocellular carcinoma (HCC) is a significant clinical challenge, with limited therapeutic options. Anthracyclines such as doxorubicin (DOX) and idarubicin (IDA) are commonly used in cancer treatment but have shown variable efficacy and side effects in HCC. This study aimed to compare the cytotoxic effects of DOX and IDA on HCC in both *in vivo* and *in vitro* models, assess their impact on the tumor and its microenvironment, as well as identify potential adverse effects. *In vivo*, both DOX and IDA treatments led to a significant reduction in body weight and spleen-to-body weight ratio, with IDA showing a more pronounced effect. However, neither treatment significantly affected intestinal morphology, compared to untreated mice with HCC. None of the treatments had a significant impact on macroscopic or microscopic tumor burden. Notably, DOX and IDA treatments resulted in a significant reduction in collagen deposition and liver fibrosis. DOX reduced hepatic stellate cell (HSC) activation, despite having no significant impact on the expression levels of fibrotic markers TGF-β and CTGF. In contrast, IDA not only increased HSC activation but also upregulated TGF-β and CTGF expression in both tumor and peritumoral tissues. Molecular analysis further revealed that DOX and IDA treatments increased mRNA levels of ER-stress and proliferation markers in non-tumor tissues, with significant findings within the PERK pathway. IDA, in particular, induced higher ATF4 expression in hepatocytes and enhanced macrophage recruitment in tissue sections. While DOX and IDA exhibit limited effectiveness in reducing HCC tumor burden *in vivo*, the *in vitro* analyses showed that DOX and IDA demonstrated strong, concentration-dependent cytotoxicity, significantly reducing cell viability in all tested HCC cell lines. Increasing complexity of the *in vitro* models, by culturing in 3D and adding HSCs, decreased the sensitivity to the anthracyclines. This, along with the effects on liver fibrosis, stellate cell activation and inflammation seen *in vivo*, may be the result of the significant contribution of the tumor microenvironment in mediating drug response. The differential expression of ER-stress and proliferation markers, particularly in the PERK pathway, further highlights the complexity of the tumor microenvironment’s influence on treatment outcomes. More research into these molecular responses and underlying mechanisms is needed to provide insights into improving therapeutic strategies for HCC.

## 1. Introduction

Hepatocellular carcinoma (HCC) is the most common type of primary liver cancer and the third leading cause of cancer-related death globally (1). HCC occurs in a context of liver disease, caused by chronic inflammation and fibrosis (2). Despite advances in therapeutic strategies, the prognosis for HCC remains poor, largely due to the tumor’s resistance to conventional chemotherapy and the complexity of its microenvironment (3, 4). Moreover, HCC is usually diagnosed at intermediate or advanced stages, making potentially curative treatments no longer available (5).

Transarterial chemoembolization, TACE, is a first-line therapy for unresectable intermediate-stage HCC (6, 7). Conventional TACE (cTACE) involves the locoregional injection of a chemotherapeutic agent close to the tumor, emulsified in Lipiodol, followed by an embolic agent to induce tumor necrosis through cytotoxic and ischemic mechanisms. An alternative to cTACE is using drug-eluting beads (DEBs), which can carry higher doses of chemotherapeutic drugs compared with cTACE and allow a gradual release (8). Despite these theoretical advantages, there are no established differences regarding safety and effect (8, 9). Anthracyclines have been widely used in cancer treatment since their discovery in the 1960s, due to their ability to intercalate DNA and inhibit topoisomerase II (10, 11). Although doxorubicin (DOX) remains a key agent in TACE, its use is associated with significant adverse effects and variability in outcomes (9, 12, 13). Therefore, there is a pressing need to find alternative therapeutic agents that can enhance efficacy while minimizing off-target toxicity.

Idarubicin (IDA), a semi-synthetic derivative of daunorubicin, is characterized by the absence of a 4-methoxy group that increases its lipophilicity and cellular uptake, suggesting a stronger cytotoxic effect against cancer cells (14). Although IDA has shown a more potent cytotoxic *in vitro* effect on different cancer models (15–18), it has also demonstrated variable responses *in vivo* (19–22). This highlights the need for an improved understanding of its mechanisms of action and the influence of tumor microenvironmental factors on the therapeutic response and resistance development.

The tumor microenvironment plays an important role in HCC progression, therapy resistance, and metastasis. It is a complex network of non-tumor cells, extracellular matrix components, and signaling molecules that interact with cancer cells, influencing their behavior and response to treatment (23). Recent studies have begun to identify the impact of the tumor microenvironment on the efficacy of chemotherapeutic agents, suggesting that the interaction between cancer cells and their microenvironment may alter drug responsiveness (24–27).

Chemotherapy is associated with several systemic adverse effects, including those affecting the gastrointestinal (GI) tract (9). Agents such as DOX and IDA specifically target rapidly dividing cells, including those of the intestinal epithelium (28, 29). The intestinal epithelium undergoes continuous renewal, making it particularly susceptible to chemotherapy-induced mucosal injury (30). Therefore, these agents can compromise intestinal integrity, resulting in epithelial atrophy, villus and crypt shortening, and impaired regenerative capacity (31). Additionally, chemotherapy-induced disruption of the gut microbiome and mucosal immune responses further exacerbates GI complications. Altogether, this can manifest in symptoms such as diarrhea, malabsorption, and weight loss. These side effects not only compromise patient quality of life, but also often necessitate treatment modifications that may reduce the overall effectiveness of HCC therapy (9). Understanding the cellular and molecular changes underlying chemotherapy-induced intestinal injury is needed to develop supportive interventions to mitigate these effects, while maintaining the anticancer efficacy of treatment.

This study aims to define the comparative effects of DOX and IDA in HCC treatment, focusing on tumor response, the tumor microenvironment and off-target effects in the intestine. We investigate their cytotoxic effects in both *in vivo* and *in vitro* models, examine their impact on fibrosis - a hallmark of HCC progression and a major component of the tumor microenvironment - explore the molecular pathways that may mediate their effects and investigate the effect on the small intestine. Through this approach, we aim to increase the understanding of how IDA and DOX affect HCC, with the goal of finding potential strategies to enhance therapeutic efficacy and overcome drug resistance.

## 2. Materials and methods

### Chemicals

Chemicals used during experiments were diethylnitrosamine (DEN; 1002877809, Sigma-Aldrich, Darmstadt, Germany, diluted in saline), doxorubicin (Doxorubicin Accord 2 mg/mL; Vnr 189790; diluted in saline), idarubicin (Zavedos®; Pfizer; Vnr 08 08 20; diluted in saline or DMSO).

### Animal model

A chemically-induced HCC murine *in vivo* model was used as previously described (32, 33). Briefly, 5-week-old male sv129-mice (129S2/SvPasCr) received intraperitoneal injections of DEN (35 mg/kg bodyweight) for 28 weeks every other week. From week 25 (when tumors have formed), mice were treated twice per week with DOX intravenous (i.v.) injections (2 µg/g bodyweight) or IDA i.v. injections (0,4 µg/g bodyweight). Control group was injected i.v. with saline. After 28 weeks, the animals were euthanized, and tissue samples were collected for analysis. Body weight was recorded in parallel with injections and at endpoint. At the end of the study, macroscopic tumors in the liver were counted and distal organs were examined for metastasis. Our protocol adhered to the Uppsala ethical committee’s standards for animal experimentation (DNR 5.8.18 0089/2020) and followed RESIST guidelines.

### Antibodies

Primary antibodies used for immunohistochemistry were Ki67 (ab16667; Abcam; 1:200), PCNA (PA5-27214; Thermo Fisher 1:100) and α-SMA (ab5694; Abcam; 1:100 and Alcian Blue-PAS (pH 2.5) (J60122.09; Thermo Fisher). Primary antibodies used for immunofluorescence were ATF4 (PA5-27576; Invitrogen; 1:100), F4/80 [CI:A3-1] (ab6640; Abcam; 1:100). Secondary antibodies used for immunofluorescence were donkey anti-rabbit Alexa Fluor™ 488 (A-21206; Invitrogen; 1:500) and donkey anti-goat Alexa Fluor™ 555 (A-21432; Invitrogen; 1:500).

### Histological staining and immunocytochemistry

Tissue blocks were prepared by 24 hours fixation in 4% paraformaldehyde, followed by paraffin embedding. Tissue sections (5 μm) were cut with a microtome and dried overnight. After deparaffinization and rehydration, slides were stained with hematoxylin and eosin (H&E) or Sirius Red (SR) following standard procedures. Immunocytochemistry was performed using the rabbit specific HRP/DAB detection IHC kit (ab64261; Abcam), following manufacturer’s instructions. Antigen retrieval was performed by 1,5-hour incubation in a decloaking chamber with DIVA decloaker 10X (DV2004MX; Biocare Medical) and primary antibodies were incubated for 1 hour at 37°C. Hematoxylin was used as counterstaining. Microscopic images were acquired with the Axio Scope Carl Zeiss microscope (10x/0.25) using the ZEN 3.2 microscopy software from ZEISS.

### Immunofluorescence

Tissue sections (5 μm) from paraffin-embedded blocks were deparaffinized and re-hydrated. Antigen retrieval was performed by 1,5-hour incubation in a decloaking chamber with DIVA decloaker 10X (DV2004MX; Biocare Medical). Tissue sections were then blocked for 1 hour with 1% BSA in PBS-Tween 20 and incubated with primary antibody in blocking buffer overnight. The next day, tissue sections were incubated with secondary antibodies in blocking buffer for 45 minutes at RT protected from light. Nuclei were stained with NucBlue™ (Hoechst 33342; R37605; Invitrogen; 1:1000). Fluorescent images were taken with a confocal laser scanning microscope (Carl Zeiss LSM 700 Laser Scanning Microscope, Jena, Germany) using a Plan Apochromat (20x/0,8) M27 Zeiss objective.

### Image analysis

Image analysis was performed with Fiji/ImageJ. For the H&E staining analysis, tumors were identified in the images and tumor area was measured. SR-stained images were analyzed by using the METAVIR score system. Furthermore, quantification of collagen deposition was performed blindly by conversion to grayscale images for each channel and automated detection of staining on thresholded images using a modified macro. α-SMA staining was analyzed by first applying a median filter with a radius of 4, followed by color deconvolution to isolate the DAB stain (34). The isolated DAB stain image was then thresholded, and the regions of interest were measured. Ki67 staining was analyzed in a similar fashion, with filter radius of 3 and particles of a determined size (70 – infinity) were quantified after thresholding. Fluorescent images were quantified by splitting the channels and measuring positive area after thresholding. Background signal was subtracted from raw values, and green (ATF4) and red (F4/80) were normalized to blue (NucBlue) positive area. To evaluate morphology and intestinal damage, villus height and width, as well as crypt depth were measured from H&E-stained samples using a straight line from the widest part of three sections per sample (35, 36). For PCNA, quantification of positive area was performed by color deconvolution and thresholding images (particle size 0-100), then taking a ratio of positively stained area to total area in sample. To quantify goblet cell area, Alcian Blue was performed similarly, instead manually counting each positively stained cell and presented as percent area of total tissue.

### Cell culture

The human liver cancer cell lines used were hepatoblastoma-derived HepG2 (ATCC. HB-8065™) and HCC-derived Huh7 (kindly gifted from Mårten Fryknäs, Uppsala University, Sweden), along with the hepatic stellate cell line LX-2 (SCC064, Sigma-Aldrich, Darmstadt, Germany) All three cell lines were routinely cultured in DMEM, high glucose, GlutaMAX™ Supplement, pyruvate (10569010; Thermo Fisher), supplemented with 1% penicillin/streptomycin (15140122; Thermo Fisher) and 10% Fetal Bovine Serum (10270106; Thermo Fisher). Starvation medium (DMEM) was only supplemented with 1% penicillin/streptomycin. Culture medium was replaced every 2-3 days and cells were split at about 70-90% confluency. Cells were maintained in an incubator at 37°C, 5% CO2 and 21% O_2_. Cell lines were authenticated and regularly tested for mycoplasma contamination.

### Monolayer viability assays

Single cells (1 x 10^4^ in 200 µL/well) were seeded in 96-well plates and allowed to attach for 24 h. To synchronize cell cycle, seeding medium was changed to serum-free (starvation) medium for overnight incubation. Stock solutions of DOX (1000 µM), and IDA (2 µM or 1000 µM), were prepared in supplemented medium and serially diluted 1:2 in fresh medium, achieving 10 concentrations, including one untreated. Starvation medium was removed and 150 µL of drug-containing medium was added per well. Treated plates were placed in the incubator for 24 h. Viability was measured by adding a 1% resazurin sodium salt solution in 1/80 dilution to the cells and incubating for 24 h (37). Fluorescent signal was measured with a 485/35 excitation filter and a 550/20 emission filter on a Fluostar Omega plate reader.

### Spheroid viability assays

For the monoculture assays, single cell solutions (2 x 10^3^ cells in 100 µL/well) were seeded in Nunclon™ Sphera™ 96-Well, Nunclon Sphera-Treated, U-Shaped-Bottom Micro-plate to generate spheroids (174925; Thermo Fisher). For the co-culture assays, single cell solutions were prepared and a ratio 4:1 of Huh7/HepG2 and LX-2 respectively were seeded per well. After the formation of spheroids in the incubator (72 h), fresh starvation medium (100 µL/well) was added and plates were incubated for 4 hours at 37°C. Stock solutions of DOX and IDA were prepared in supplemented medium at 2X final concentration and serially diluted 1:2 in fresh medium. Next, fresh medium (100 µL) containing drug was added to each well resulting in a total volume of 200 µL and achieving a 1X drug concentration. Spheroids were treated with DOX (0 to 1000 µM), IDA (0 to 1000 µM) or IDA (0 to 1000 µM) + AMG-PERK44 (10 µM). Treated spheroids were placed in the incubator for 24 hours. After 24 hours, viability was measured using the CellTiter-Glo® 3D Cell Viability ATP-based assay (G9683; Promega), following manufacturer’s instructions. Luminescence was recorded with the FLUOstar Omega plate reader.

### Quantitative RT-PCR of mRNA

Total RNA was extracted using the E.Z.N.A. Total RNA-Kit I (R6834-02, Omega Bio-tek), following manufacturer’s guidelines. RNA yield was measured using Nanodrop. cDNA was obtained using the iScript cDNA-synthesis kit (1708891, Bio-rad), following manufacturer’s guidelines. Primers were designed with NCBÍs Primer-BLAST and ordered from ThermoFisher (Supplementary Material Table 1). qPCR was performed in duplicates in a 384-well plate using Fast SYBR Green Master Mix (434385612; Thermo Fisher), according to manufacturer’s instructions. QuantStudio 5 (ThermoFisher Scientific) was used to analyze the data, and fold change was calculated using the 2-ΔΔ Ct method. Target gene expression levels were normalized to the reference genes (GAPDH and α-actin) and relative expression levels were further normalized to healthy control group.

### Statistical analysis

Data are presented as mean ± standard error of the mean (SEM) and p-values <0.05 were considered statistically significant. Statistical significance was determined using one-way analysis of variance (ANOVA) followed by Tukey’s multiple comparison test. For viability assays, data points are the averaged values of technical replicates from two independent biological experiments. Statistical relevance was verified using non-linear regression. For qPCR experiments, outliers were identified using the ROUT method and excluded from the analysis. Two-way ANOVA with Tukey’s multiple comparisons test was performed on cleaned data. Statistical analysis of intestinal samples was performed using an ANOVA with Tukey’s HSD post-hoc test, with comparisons of p<0.05 considered significant. When necessary, pairwise comparisons were performed using a student’s t-test. All statistical analyses and graphs were made using GraphPad Prism 9.

## 3. Results

### Effects of DOX and IDA treatment on body weight, spleen ratio, tumor burden, and cellular proliferation in a DEN-induced HCC mouse *in vivo* model

Mice with DEN-induced HCC presented with a significant reduction in body weight compared to the healthy and untreated group (Figure 1A). Treatment with DOX and IDA significantly aggravated this reduction in body weight, an expected side effect of anthracyclines (Figure 1A). This effect was more pronounced in the IDA group. IDA treatment significantly increased the spleen-to-body weight ratio in mice with DEN-induced HCC (Figure 1C). Upon evaluating the therapeutic efficacy of DOX and IDA against HCC in this model, both agents demonstrated a limited ability to diminish macroscopic tumor numbers (Figure 1B). Further histological examination substantiated that neither drug significantly mitigated the microscopic tumor burden within the liver (Figure 1D and 1F). However, treatment with IDA reduced the relative tumor area, compared to the relative tumor area quantified in the untreated DEN group (Figure 1D). Immunohistochemical staining with the proliferation marker Ki67 showed a significant increased number of Ki67 positive cells in the DEN-induced HCC group (Figure 1E and 1F), which was slightly reduced by IDA-treatment (Figure 1E).

**Figure 1:**
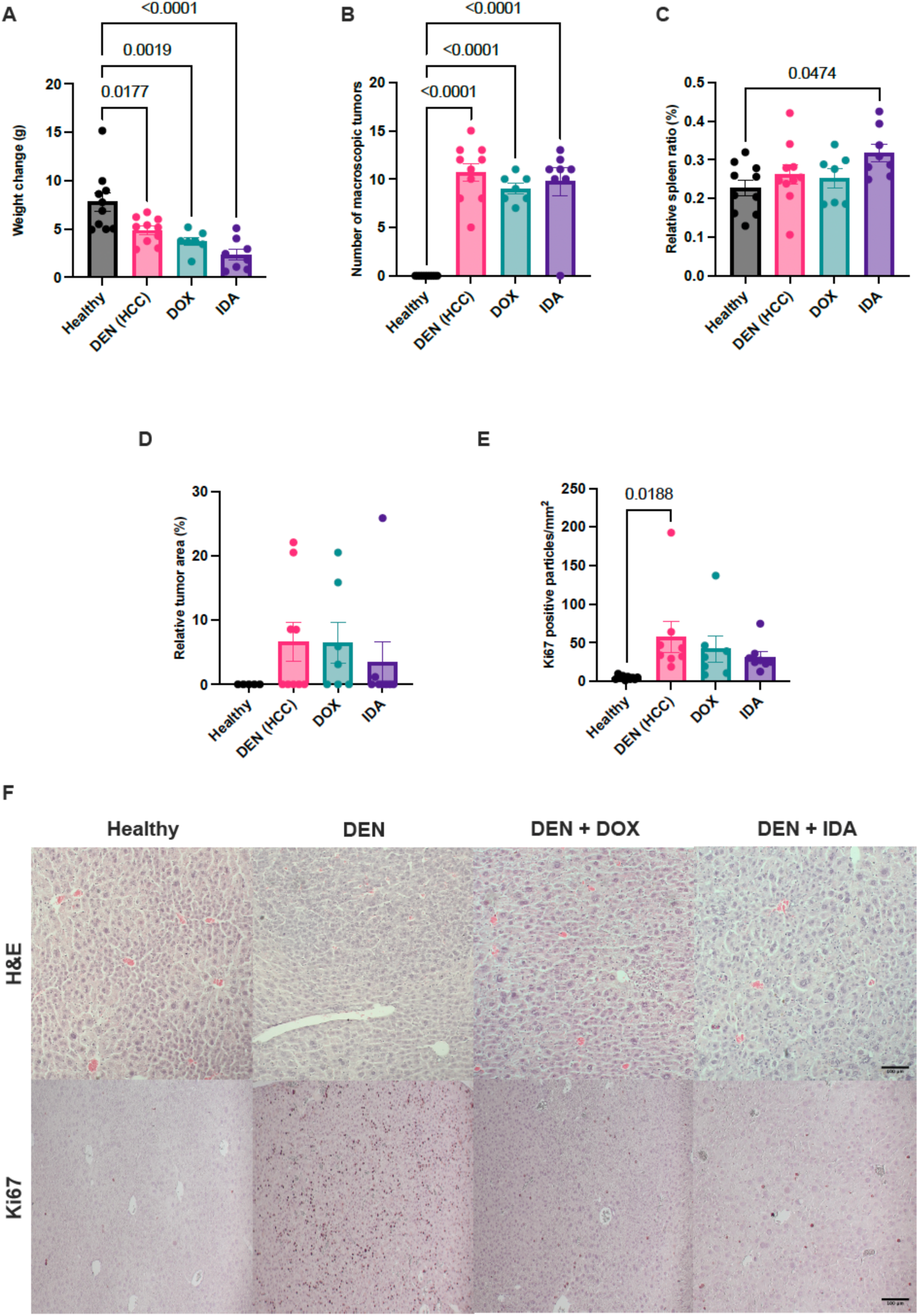
Effects of DOX and IDA on DEN-induced HCC in mice (in vivo). (A) Body weight change of healthy control mice, DEN-induced HCC mice, and DEN-induced HCC mice treated with either doxorubicin (DOX) or idarubicin (IDA). (B) Relative spleen ratio. (C) Number of macroscopic tumors (D) Quantified relative tumor area from tissue sections. (E) Quantified Ki-67 positive cells from tissue sections (F) Representative images of tissue sections stained with hematoxylin and eosin (H&E) and immunohistochemical staining for Ki-67. Each data point represents one mouse. N = 5-10 mice per group. All data is expressed as mean ± SEM. Scale bars represent 100 µm.

### Effects of DOX and IDA treatment on proliferation, **mucin** production, and intestinal morphology in a DEN-induced HCC mouse in vivo model

As we observed significant weight loss in the DOX and IDA treated mice, we wanted to establish whether this was due to chemotherapy-induced mucosal damage in the small intestine, as this is a common side-effect of anthracyclines and known contributor to anorexia in cancer patients receiving chemotherapy. However, H&E staining of jejunal sections showed no significant morphological changes in villus length across experimental groups (Figure 2A and E). This suggests that villus integrity is maintained despite chemotherapy exposure and HCC progression.

**Figure 2.**
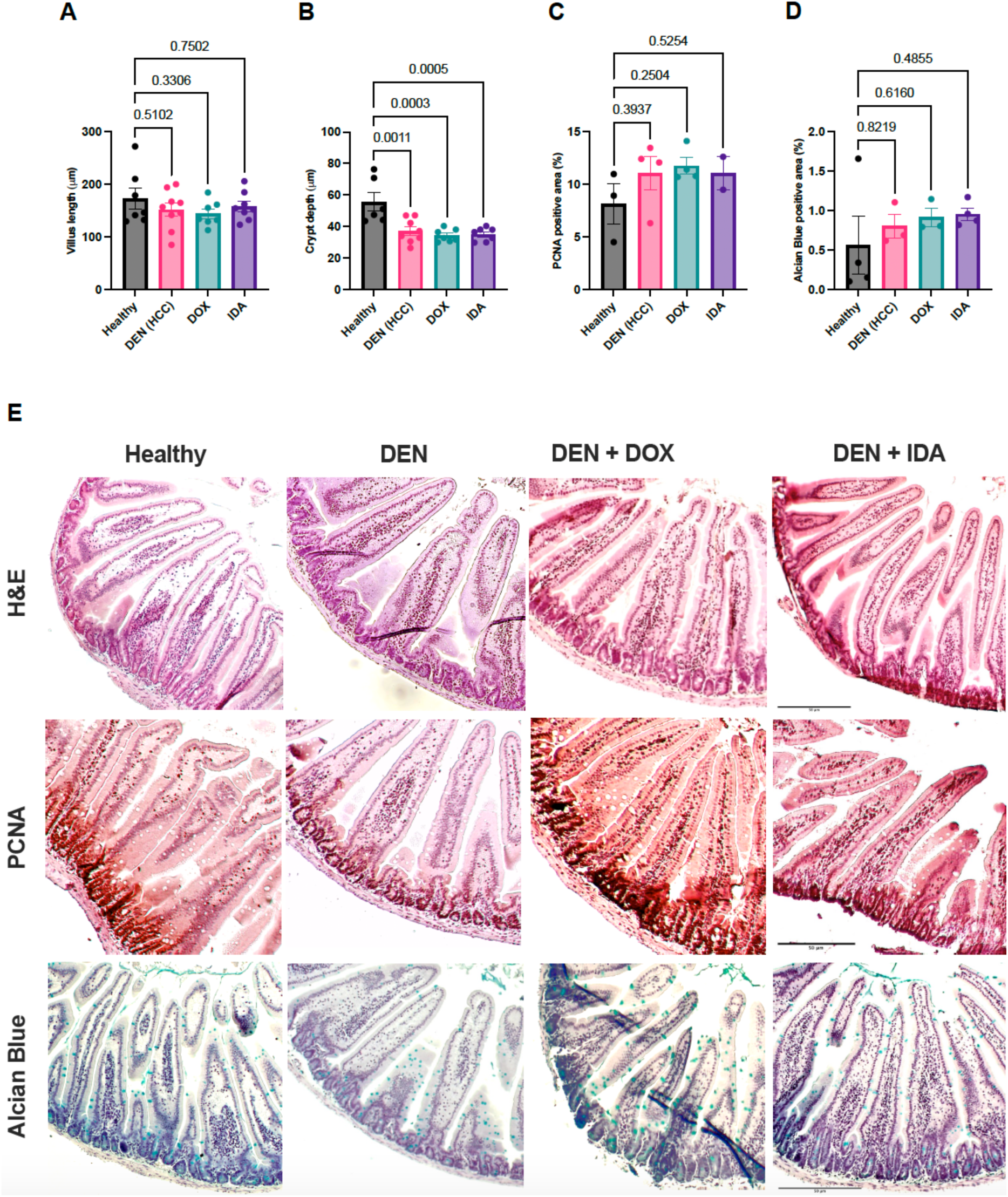
Histological and immunohistochemical analysis of intestinal sections in experimental groups. Villus length (A) and crypt depth (B) measurements of the three jejunal cross-sections were taken per mouse. Quantified PCNA (C) and Alcian Blue (D) positive area from jejunal sections. Each data point represents averaged values from one mouse. N = 5-10 mice per group. Data are presented as mean ± SEM, with statistical significance indicated where applicable. Scale bars represent 50 µm.

In contrast, crypt depth was significantly reduced across all mice with DEN-induced HCC compared to the healthy controls (Figure 2B and E). This reduction was slightly more pronounced in chemotherapy-treated mice with HCC, indicating the presence of crypt atrophy. Given that crypts house intestinal stem cells responsible for epithelial renewal, these results suggest that both the DEN-treatment and the chemotherapy suppress regenerative capacity, thereby compromising intestinal homeostasis, which aligns with previous literature (38).

To evaluate the regenerative response following chemotherapy exposure, proliferative activity was assessed via PCNA staining (Figure 2C and E). PCNA-positive staining did not significantly change in any treatment group. However, there was a trend to increased PCNA-positive area in all mice with DEN-induced HCC compared to the healthy controls. These results indicate that, despite crypt atrophy, a compensatory increase in epithelial cell proliferation was not observed.

The integrity of the mucosal barrier was further examined using Alcian Blue staining to assess mucin production by goblet cells (Figure 2D and E). No significant differences in Alcian Blue-positive staining were detected between groups. However, there was a trend towards increased number of goblet cells in all mice with DEN-induced HCC, with slightly higher levels in those receiving IDA or DOX. These results indicate that mucin production might have increased in both the DEN-treatment and after chemotherapy-induced epithelial injury.

### IDA contributes to the maintenance of an inflammatory and fibrotic microenvironment in a DEN-induced HCC mouse model

HCC develops in a context of chronic inflammation and liver fibrosis (39, 40). Furthermore, activated hepatic stellate cells (HSCs) play a major role in the pathogenesis of HCC, contributing to liver fibrosis through excessive collagen production and pro-inflammatory signaling (2, 25, 41). Treatment with DOX and IDA led to a notable reduction in collagen deposition, as shown by histological quantification of Sirius red staining on liver slides (Figure 3A and 3E). This histological evidence of reduced fibrosis was further confirmed by Metavir scoring (Figure 3B). We then analyzed the expression of α-SMA, a specific marker for HSC activation. In tissue sections, there was a significant increase in α-SMA positive area in the DEN-induced HCC group (Figure 3C and 3E). Treatment with DOX significantly reduced the α-SMA positive area relative to DEN. Conversely, the α-SMA positive area in the IDA-treated group remained similar to that observed in the DEN group. Further qPCR analysis confirmed significantly increased mRNA levels of α-SMA in the non-tumor tissue of DEN-induced HCC mice (Figure 3D). Interestingly, while mRNA levels of α-SMA were similar in the non-tumor tissue of both DEN and DOX-treated mice, treatment with IDA reduced these mRNA levels. Overall, these data suggest that while DOX and IDA exhibited limited effectiveness in reducing the tumor burden, but they significantly decreased collagen deposition in a DEN-induced HCC murine model.

**Figure 3:**
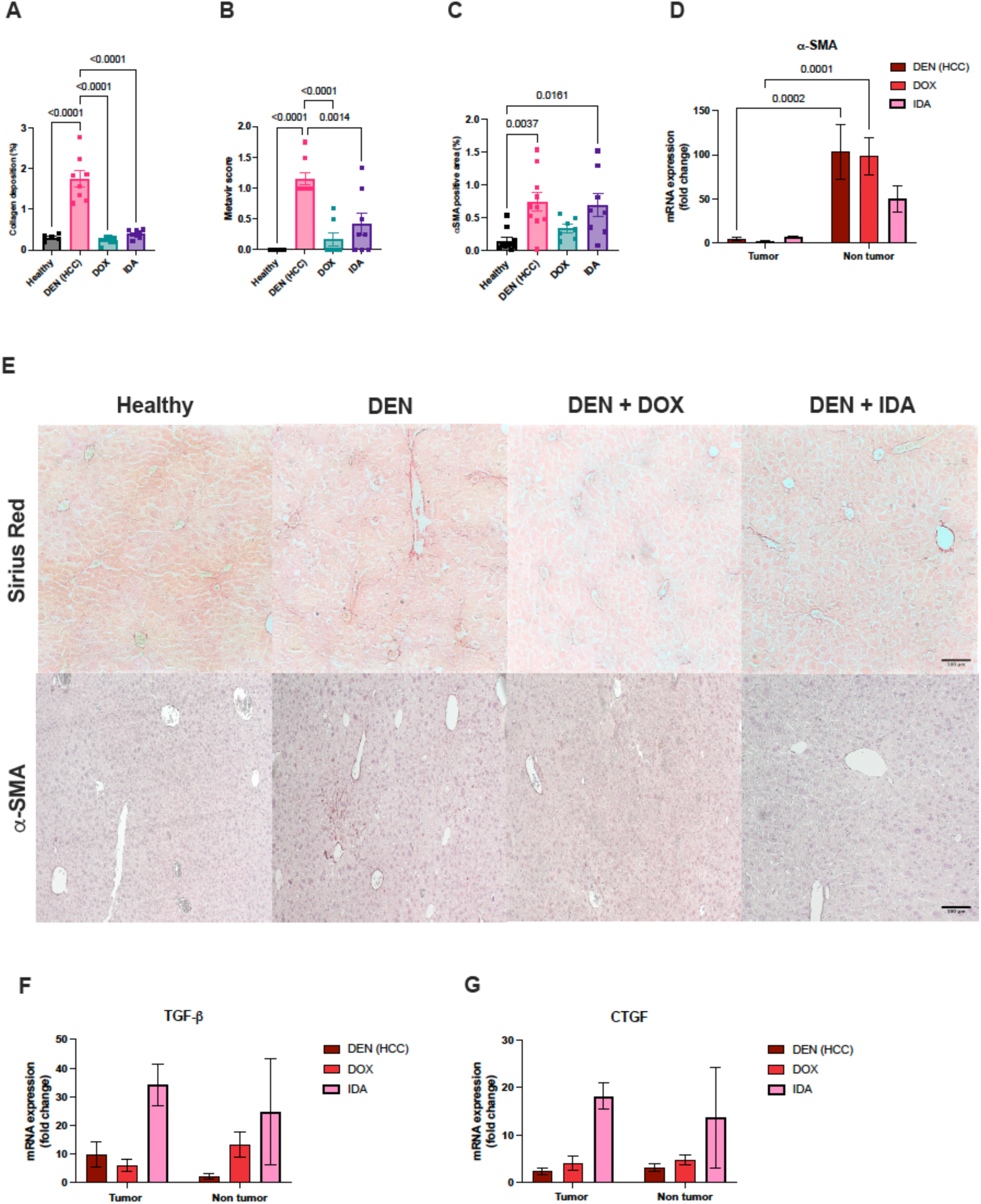
Effects of DOX and IDA on fibrosis and hepatic stellate cell activation in a murine DEN-induced HCC model. (A) Collagen deposition. The bar graph represents percentage of collagen deposition quantified from Sirius Red-stained liver slides. (B) Metavir score. (C) α-SMA staining quantification. The bar graphs show the percentage of quantified positive area from α-SMA-stained liver slides. (D) mRNA expression levels of α-SMA in murine liver tissue. (E) Representative images of Sirius Red and α-SMA-stained liver samples. mRNA expression levels of TGF-β (F) and CTGF (G) in murine liver tissue. Each data point represents one mouse. N = 5-10 mice per group. All data is expressed as mean ± SEM. Scale bars represent 100 µm.

In the context of liver disease, TGF-β is a primary cytokine that activates HSCs, leading to their transformation into fibrogenic myofibroblasts and promoting extracellular matrix (ECM) production (41–43). Connective-tissue growth factor (CTGF), induced by TGF-β, further enhances and sustains the fibrotic response by amplifying fibrogenic signals and supporting continued ECM accumulation (43). In our study, we found that both CTGF and TGF-β were upregulated in tumor and non-tumor tissue from mice treated with IDA (Figure 3F and 3G). This is in line with many findings that show that treatment with anthracyclines induces TGF-β release and signaling (44–46). Conversely, DOX only upregulated TGF-β expression in non-tumor tissue.

To quantify the effect of DOX and IDA on inflammation, mRNA-levels of four inflammatory markers were measured in tumor and non-tumor tissue of mice treated with IDA and DOX. Overall, mRNA-levels of inflammatory markers were higher in the IDA-treated group, both in tumor and non-tumor tissues (Figure 4).

**Figure 4:**
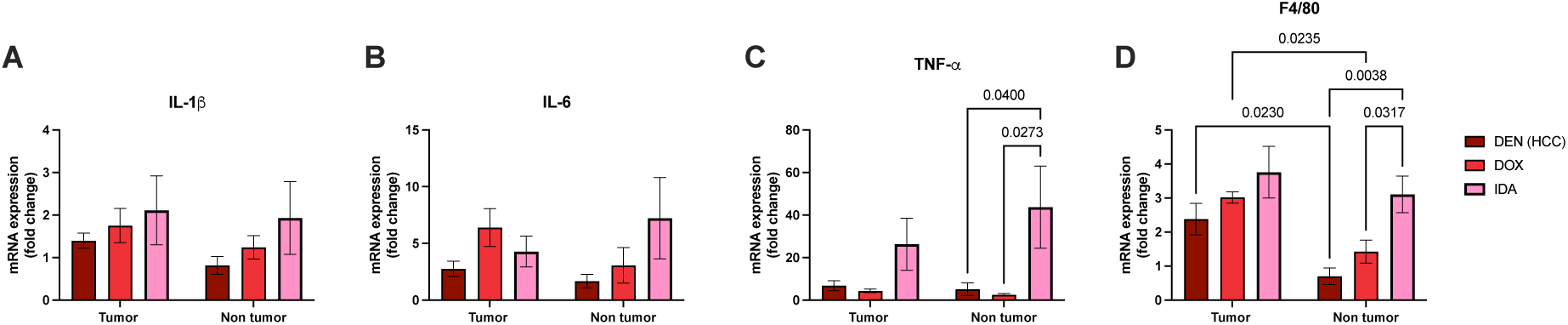
Idarubicin treatment increases mRNA expression of inflammation markers. mRNA expression levels of IL-1β (A), IL-6 (B), TNF-α (C) and F4/80 (D) in DEN-induced HCC mice, and DEN-induced HCC mice treated with either DOX or IDA. N = 5-10 mice per group. All data is expressed as mean ± SEM.

Similar levels of IL-1β expression were observed in tumor and non-tumor tissues upon treatment (Figure 4A). Interestingly, IL-6 expression was upregulated by DOX in tumor tissue and by IDA in non-tumor tissue (Figure 4B). IDA treatment significantly increased TNF-α expression in non-tumor tissue (Figure 4C). The expression of macrophage marker F4/80 was higher in tumor than in non-tumor tissues (Figure 4D). These increases in IL-6, TNF-α, and F4/80 after IDA treatment aligns with known mechanisms of chemotherapy-induced inflammation and macrophage activation, which can be both a response to and a cause of tissue injury (32).

### Treatment with IDA increases ATF4 expression in hepatocytes and recruits macrophages

To assess how these treatments affected ER-stress pathways, we analyzed the mRNA-expression of several ER-stress related genes, namely homocysteine-inducible ER-stress protein (HERP), ER oxidoreductin 1β (Ero1b), heat shock protein family A member 5 (HSPA5) and ER degradation enhancing alpha-mannosidase like protein 1 (Edem1) (Figure 5A-G). In general, DOX and IDA seemed to have similar effects on ER-stress markers; expression was generally higher in the non-tumor tissue than in the tumor tissue, which is in line with previous findings (40, 47, 48).

**Figure 5:**
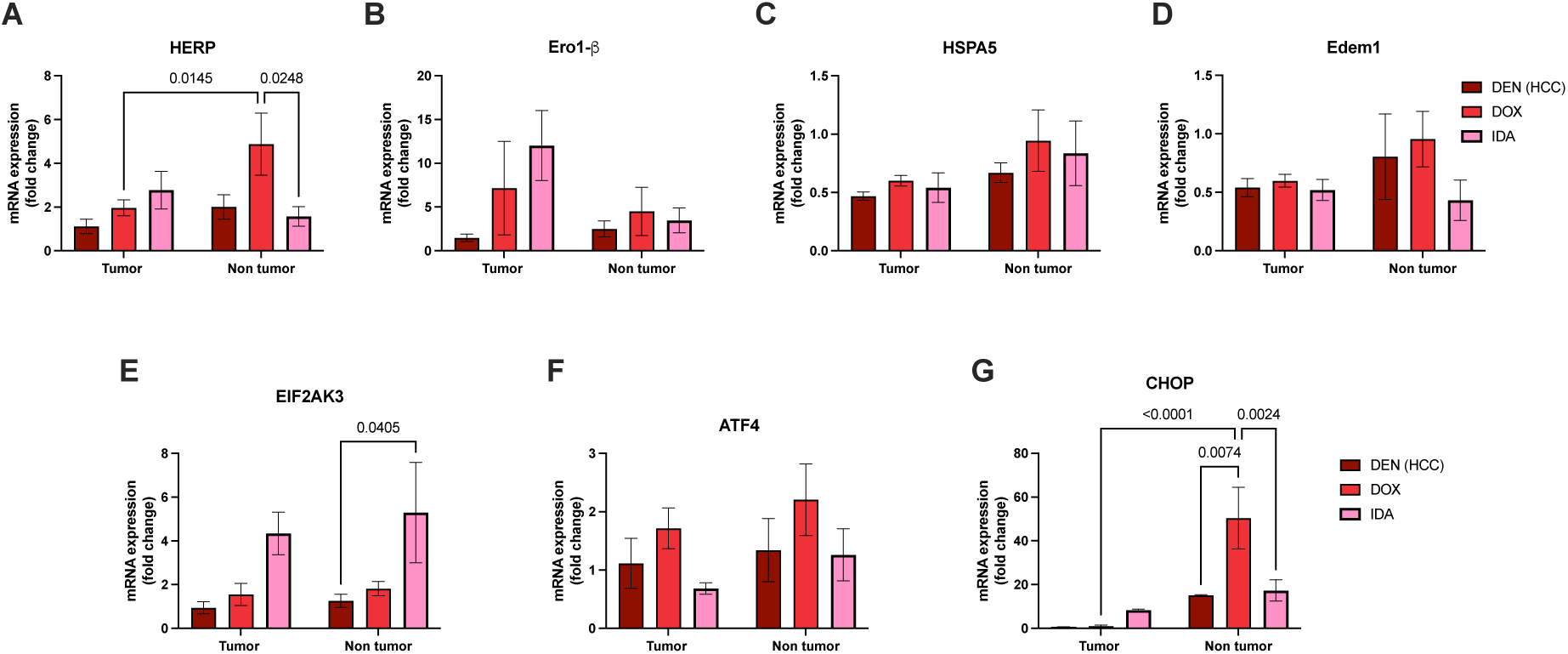
ER-stress markers qPCR and fluorescence. mRNA expression levels of HERP (A), ERO1β (B), HSPA5 (C), Edem1 (D), EIF2AK3 (E), ATF4 (F) and CHOP (G) in DEN-induced HCC mice, and DEN-induced HCC mice treated with either DOX or IDA. N = 5-10 mice per group. All data is expressed as mean ± SEM.

**Figure 6:**
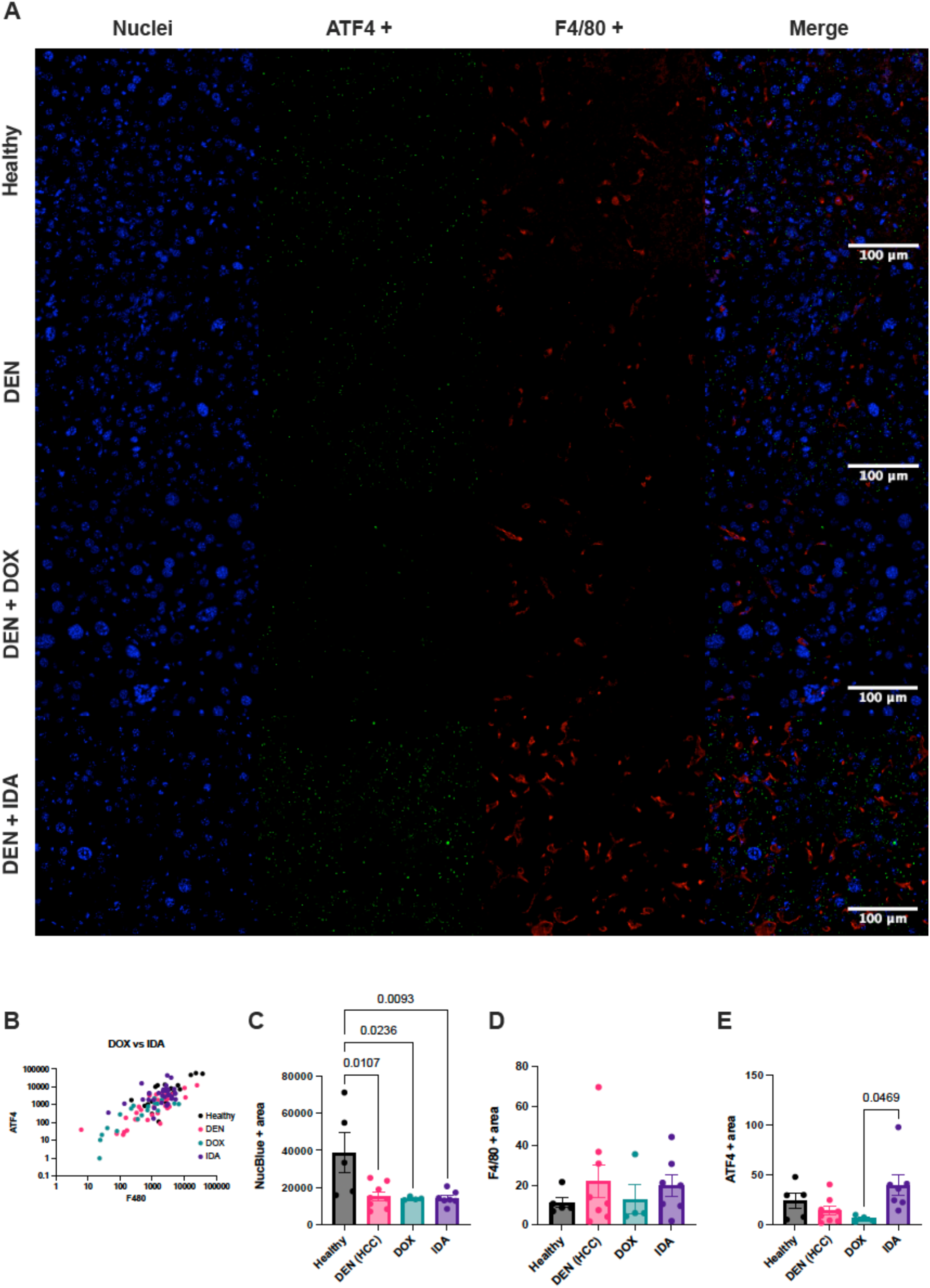
Treatment with IDA increases ATF4 expression in hepatocytes and recruits macrophages. (A) Representative images of ATF and F4/80 immunofluorescent staining on tissue sections from healthy control, DEN-induced HCC mice and DEN-induced HCC mice treated with either DOX or IDA. (B) Scatter plot showing the correlation between ATF4 and F4/80 among the groups. Quantification of nuclei (Nucblue+ area) (C), ATF4 (ATF4+ area) (D) and F4/80 (F4/80+ area) (E) signal from tissue sections. N = 5-10 mice per group. All data is expressed as mean ± SEM. Scale bars represent 100 µm.

DOX treatment significantly increased the expression of HERP in non-tumor tissues (Figure 5A). Conversely, IDA and DOX increased ERO1-β expression in tumor tissues (Figure 5B). HSPA5 and Edem expression levels remained relatively similar across all conditions in both tumor and non-tumor tissues, with non-tumor tissues showing slightly higher expression overall (Figure 5C, 5D). Notably, protein kinase RNA-like endoplasmic reticulum kinase (EIF2AK3) expression was substantially higher in non-tumor tissue of the IDA-treated group (Figure 5E). Activating transcription factor 4 (ATF4) showed a similar pattern in both tumor and non-tumor tissues; increased expression with DOX and decreased expression with IDA (Figure 5F). Interestingly, C/EBP Homologous Protein (CHOP) expression was significantly higher in non-tumor tissue of the DOX-treated group (Figure 5G). These observations underscore the nuanced influence of IDA and DOX on ER-stress responses and cellular stress mechanisms, reflecting a differential modulation of these pathways in tumor versus non-tumor liver tissues.

We found higher mRNA levels of ATF4 in the non-tumor tissue (Figure 5F). On the other hand, higher mRNA levels of macrophage marker were found on the tumor tissue (Figure 4D). We then decided to further examine and quantify these markers at the protein level on tissue sections. Fluorescent staining revealed that there was no co-localization of ATF4 and F4/80 in the tissue sections (Figure 6A). While ATF4 signal was found on the hepatocytes, F4/80 signal was found in the areas surrounding the hepatocytes. However, although there was no co-localization, the scatter plot revealed a strong positive linear correlation between ATF4 and F4/80 in all the groups (Figure 6B). Further quantification of the positive signal indicated that ATF4 and F4/80 expression was significantly increased in the IDA-treated group, while a decrease was seen in the DOX-treated group (Figure 6D and 6E).

**Figure 6:**
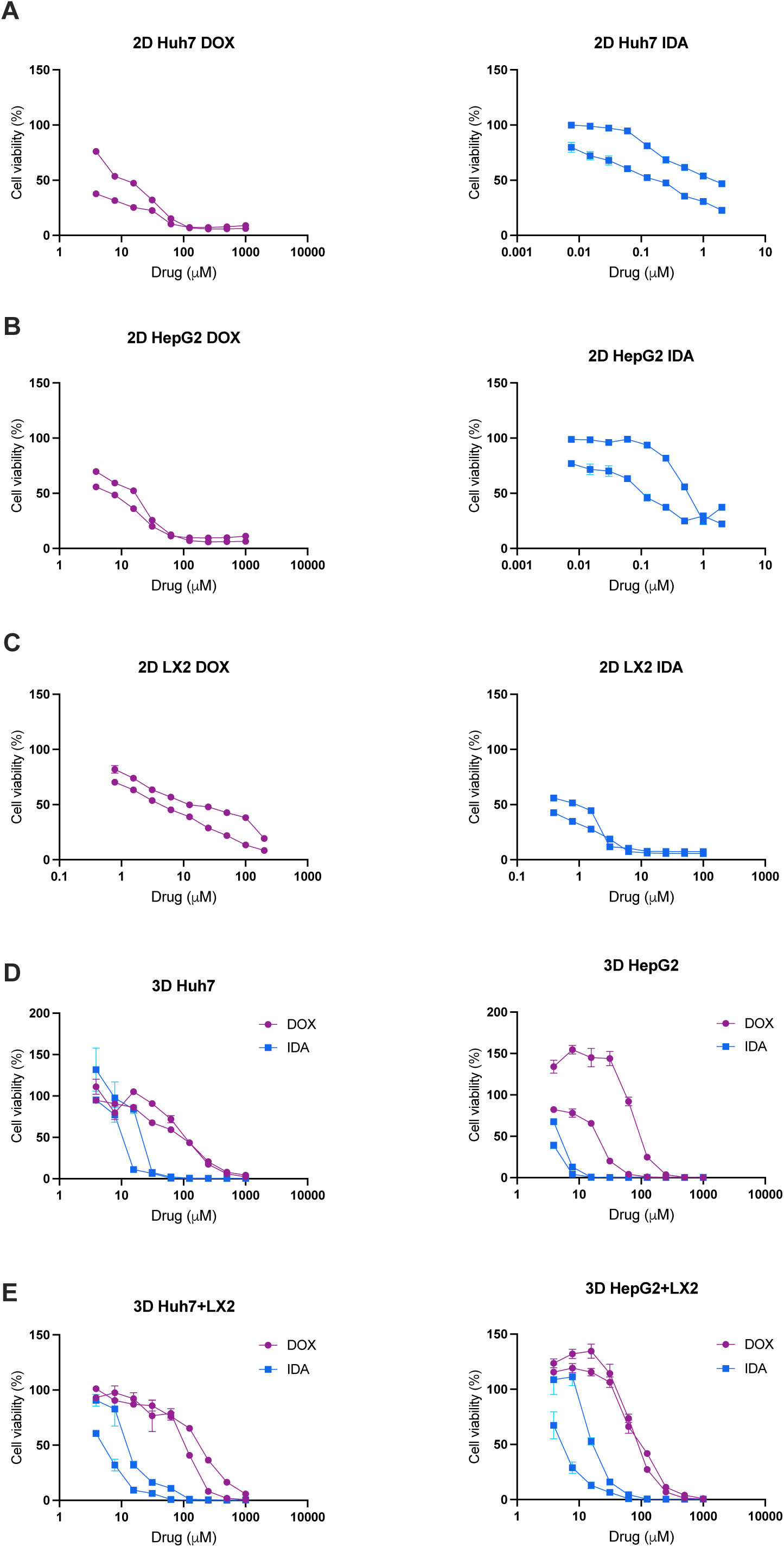
Treatment with DOX and IDA reduces cell viability on HCC cell lines *in vitro*. Cell viability curves of HCC cell lines (Huh7 and HepG2) treated with anthracyclines Doxorubicin and Idarubicin for 24 hours (A-F). Cell viability curves of Huh7 (A), HepG2 (B) and LX-2 (C) cells treated with DOX and IDA on monolayer. (D) Cell viability curves of Huh7 and HepG2 spheroids treated with DOX and IDA. (E) Cell viability curves of Huh7-LX-2 and HepG2-LX-2 spheroids treated with DOX and IDA. Each data point represents the averaged values of six technical replicates from one independent experiment. N = 2 independent experiment. Data is expressed as mean ± SEM.

### Cytotoxicity of IDA and DOX is affected by 3D culture conditions and by co-culturing with hepatic stellate cells

To assess the cytotoxic effects of IDA and DOX in different cell culture models, we used Huh7 and HepG2 cell lines grown in both 2D and 3D cultures. Additionally, 3D co-cultures with hepatic stellate cells (LX-2) were used to simulate the tumor microenvironment and evaluate its influence on drug response. In 2D cultures, IDA showed significantly higher potency compared to DOX (Figure 7A and 7B). Specifically, in Huh7 cells, IDA was 36 times more potent than DOX, with IC50 values of 0.15 µM for IDA and 5.6 µM for DOX (Table 1). Similarly, in HepG2 cells, IDA was 30 times more potent than DOX, with IC50 values of 0.26 µM for IDA and 8.0 µM for DOX. Interestingly, LX-2 cells alone were more sensitive to DOX and more resistant to IDA than Huh7 and HepG2 (Table 1).

**Table 1:**
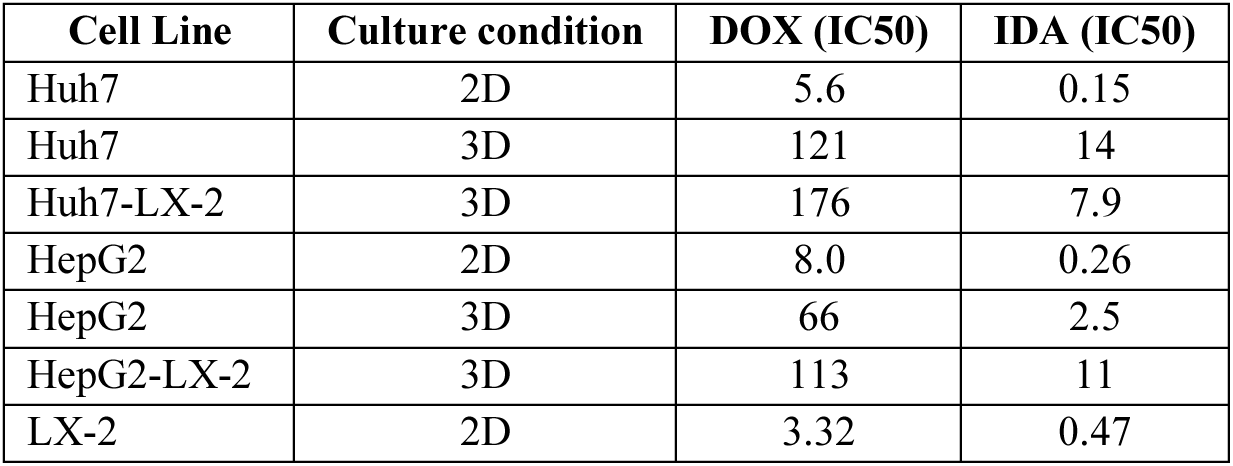
IC50 values (µM) of idarubicin (IDA) and doxorubicin (DOX) in different cell cultures.

The transition from 2D to 3D spheroid culture resulted in decreased sensitivity to both drugs. In Huh7 cells, 21 times more DOX was required to achieve 50% cell death in 3D culture (IC50 = 121 µM) compared to 2D culture (IC50 = 5.6 µM) (Table 1). For HepG2 cells, an 8-fold increase in DOX concentration was needed in 3D culture (IC50 = 66 µM) compared to 2D culture (IC50 = 8.0 µM). This decreased sensitivity was even more pronounced for IDA. In Huh7 cells, a 94-fold increase in IDA concentration was necessary to achieve 50% cell killing in 3D culture (IC50 = 14 µM) compared to 2D culture (IC50 = 0.15 µM) (Table 1). For HepG2 cells, a 9-fold increase in IDA concentration was required in 3D culture (IC50 = 2.5 µM) compared to 2D culture (IC50 = 0.26 µM).

To investigate the impact of the tumor microenvironment on drug response, we included hepatic stellate cells (LX-2) in the 3D spheroid cultures (Figure 7D). In Huh7-LX-2 3D co-cultures, the sensitivity to DOX slightly decreased, with a 1.45-fold increase in the drug concentration needed to achieve 50% reduction in cell viability (IC50 = 176 µM) compared to Huh7 3D monocultures (IC50 = 121 µM) (Table 1). Interestingly, for IDA, the co-culture condition required slightly less drug to kill 50% of the cells (IC50 = 7.9 µM) compared to the Huh7 3D monoculture (IC50 = 14 µM). In HepG2-LX-2 3D co-cultures, both DOX and IDA showed decreased sensitivity. The IC50 for DOX increased by 1.7-fold (IC50 = 113 µM) compared to HepG2 3D monocultures (IC50 = 66 µM), and for IDA, a 4-fold increase was observed (IC50 = 11 µM) compared to HepG2 3D monocultures (IC50 = 2.5 µM) (Table 1).

## 4. Discussion

Doxorubicin remains a key agent in TACE treatment for unresectable intermediate-stage HCC (6, 7). However, its use is associated with significant adverse effects and variability in outcomes (6, 7, 11–13). In addition, there is a heterogeneity of applied TACE treatment approaches that also vary extensively among the different global regions, hospitals, and even colleagues within the same interventional radiologist team (6, 7). Therefore, there is a need to investigate alternative therapeutic agents to enhance efficacy while minimizing toxicity. One such agent idarubicin (IDA), a semi-synthetic derivative of daunorubicin, characterized by the absence of a 4-methoxy group, increasing its lipophilicity and cellular uptake (14). IDA has shown a more potent cytotoxic *in vitro* effect on different cancer models (15–18), but there have been variable responses *in vivo* (19–22). This underscores the need for further research to evaluate whether idarubicin could serve as a safer and more effective alternative to DOX in the treatment of unresectable intermediate-stage HCC.

In our *in vivo* animal study, both DOX and IDA showed limited effectiveness in reducing the number of macroscopic HCC tumors in mice with DEN-induced HCC. Histological analyses confirmed that neither DOX nor IDA significantly decreased the microscopic tumor burden, with also no discernible differences in the number of Ki67-positive cells. This aligns with previous findings indicating that DEN-induced liver tumors share similarities with HCC in patients with low response to chemotherapy and poor prognosis (49, 50). Both DOX and IDA treatments were associated with increased body weight loss, an expected side effect of anthracyclines (51). Intriguingly, IDA treatment increased the spleen-to-bodyweight ratio in mice with DEN-induced HCC. The observed increase in spleen size (splenomegaly) in mice with DEN-induced HCC treated with IDA could have multiple explanations beyond portal hypertension. One potential factor is the immunomodulatory effects of the anthracyclines (52, 53). Anthracyclines like IDA may influence immune cell populations, potentially leading to alterations in spleen size due to changes in immune cell proliferation and/or infiltration (53–55). Additionally, the spleen plays a key role in filtering damaged or senescent red blood cells; IDA-induced toxicity could result in increased splenic workload and subsequent enlargement; or as a result of direct injury to the spleen (51, 56–58). Therefore, the splenomegaly observed may result from a combination of factors, including portal hypertension, immune modulation, multi-organ toxicity and hematologic effects associated with IDA treatment. As both anthracyclines led to a significant reduction in the Metavir score of Sirius red-stained liver slides, indicating a decrease in fibrosis, this would likely eliminate the portal hypertension hypothesis.

HCC develops in a context of chronic inflammation and liver fibrosis (39, 40). Furthermore, activated HSCs play a major role in the pathogenesis of HCC, contributing to liver fibrosis through excessive collagen production and pro-inflammatory signaling (43). The findings in this study demonstrate that DOX and IDA significantly decreased collagen deposition in a DEN-induced HCC murine *in vivo* model. As evidenced in our research, DOX and IDA demonstrated distinct and unexpected effects on HSC activation and fibrotic signaling. DOX notably reduced HSC activation, despite having no significant impact on the expression levels of key fibrotic markers such as TGF-β and CTGF, which remained comparable to those in the untreated DEN group. In contrast, IDA not only increased HSC activation but also markedly upregulated TGF-β and CTGF expression in both tumor and peritumoral tissues. These contrasting effects highlight a surprising dichotomy in the way DOX and IDA influence the fibrotic microenvironment, with IDA promoting a profibrotic response that aligns with its effect on HSC activation, while DOX appears to decouple HSC activation from fibrotic marker expression. TGF-β is a well-known driver of fibrosis and tumor progression, promoting the synthesis of ECM-components and facilitating epithelial-to-mesenchymal transition (EMT) (59, 60). The upregulation of CTGF, which acts downstream of TGF-β, further supports the fibrotic response and tissue remodeling within the tumor microenvironment. This could enhance tumor growth and invasiveness. Anthracyclines could potentially induce a stress response in tumor cells (61), leading to increased secretion of fibrogenic factors, like TGF-β and CTGF (62), to support tumor survival and adaptation under chemotherapeutic pressure (63).

The observed reduction of α-SMA protein and collagen deposition suggests that DOX might inhibit myofibroblast activation and/or collagen synthesis. This could be due to a differential response of non-tumor tissue to the chemotherapeutic agents, where the (dividing) non-tumor cells might be more susceptible to the cytotoxic effects, leading to reduced myofibroblast activity and ECM production. Recent studies have shown that DOX can induce autophagy in fibroblasts, which could have contributed to the removal of stellate cells, despite the increased release of TGF-β in the tumor compartment (64). Others studies have also shown that DOX can modulate the TGF-β signaling cascade in mouse fibroblasts, inhibiting the TGF-β induced expression of collagen in mouse embryonic fibroblasts (46). TGF-β signaling has widely been implicated in anthracycline toxicity, vastly contributing to its adverse effects on non-tumor tissues (44–46) and contributing to therapeutic resistance (63).

The increase in IL-6, TNF-α, and F4/80 after idarubicin treatment aligns with known mechanisms of chemotherapy-induced inflammation. This inflammatory response contributes to HSC activation and reflects the well-documented mechanisms of chemotherapy-induced inflammation, particularly with anthracyclines (65). The observed cytokine increase and macrophage activation indicate an immune response, which could be a result of the oxidative stress and/or cellular damage induced by idarubicin (44, 56, 64). Chemotherapeutics, including anthracyclines, are known to promote macrophage infiltration and create an inflammatory environment (53), which exacerbates tissue injury and highlights the dual role of macrophage activation as both a driver and responder to chemotherapy-induced damage (66). As the inflammatory response and activation of ER stress pathways are highly intertwined (30, 40, 47, 67), we quantified expression of ER-stress markers in tumor and non-tumor tissue of mice with DEN-induced HCC treated with IDA and DOX. In spite of the differential expression patterns observed for DOX and IDA, treatment with anthracyclines lead to a generally higher expression of ER-related genes in the peritumoral tissue than in the tumoral tissue, which is in line with previous findings (40, 47, 48). To further investigate the link between the inflammatory response and the ER-stress, we assessed co-localisation of ATF4 and F4/80 on tissue sections. Fluorescent staining revealed that indeed, there was no co-localization of ATF4 and F4/80 in the tissue sections. However, there was a correlation where high levels of ATF4 in hepatocytes were correlated to high levels of macrophage infiltration. This could indicate that the anthracyclines induce ER-stress in the hepatocytes, leading to activation of pro-apoptotic pathways and triggering an inflammatory response, causing macrophage recruitment.

IDA has been reported to be 5 times more potent than DOX on different cancer cell lines *in vitro* (14, 15, 18). To further evaluate the cytotoxic effect of DOX and IDA on different *in vitro* cell culture models, we performed cytotoxicity assays on Huh7 and HepG2 cell lines grown in both 2D and 3D cultures. Additionally, 3D co-cultures with hepatic stellate cells (LX-2) were used to simulate the tumor microenvironment and further evaluate its influence on drug response. As reported in the literature, IDA demonstrated to be 7 to 36 times more potent than DOX in all culture conditions. Furthermore, both Huh7 and HepG2 cell lines exhibited significantly increased resistance to both DOX and IDA when cultured in 3D compared to 2D. This supports the increasing evidence that 3D cultures better mimic the *in vivo* environment, often resulting in requiring higher drug load. Interestingly, LX-2 cells exhibited a differential drug sensitivity profile, being more sensitive to DOX and more resistant to IDA compared to Huh7 and HepG2. This increased sensitivity of LX-2 cells to DOX supports our hypothesis that DOX exerts its effects by selectively targeting and eliminating hepatic stellate cells, which could contribute to the acute inflammatory response observed in our DEN-induced HCC mouse model. Nevertheless, the inclusion of LX-2 in 3D co-cultures further reduced drug sensitivity, suggesting complex interaction between tumor cells and stellate cells that can differentially affect drug efficacy. These observations underscore the importance of using advanced cell culture *in vitro* models to better understand the complexities of drug response and resistance, as well as increase the possibilities for a successful translation to clinical settings.

This study also investigated the impact of chemotherapy and DEN-induced HCC on intestinal injury using histological and immunohistochemical techniques. Both the DEN-treatment and the chemotherapy significantly reduced crypt depth, indicating intestinal atrophy, likely through suppression of stem cell-driven epithelial regeneration. Interestingly, while chemotherapy did not notably affect villus length, proliferation, or mucin production, there was a slight trend toward increased proliferation (PCNA staining) that did not reach significance, suggesting potential disruptions in regenerative signaling or delayed proliferation at a different time point. This observation calls for further mechanistic investigation into stem cell depletion or functional inhibition. Of note, Alcian Blue staining showed no significant changes in goblet cell numbers, but again a trend was seen towards higher levels of goblet cells in the DEN-treated and chemotherapy treated mice. Importantly, our findings also highlighted that DEN-treatment alone caused intestinal damage, consistent with earlier studies linking DEN to oxidative stress and inflammation in the colon. The combination of DEN and chemotherapy treatment enhanced the reduction in crypt depth, a rarely investigated effect that calls for further exploration, especially considering potential cautions when using this mouse model in future studies. A study investigating early destructive changes caused by DEN as reported significant oxidative stress and inflammation in the colon of treated rats. Specifically, these were marked by an increase in malondialdehyde, a lipid peroxidation marker, and pro-inflammatory cytokines such as interleukin-1 and tumor necrosis factor. Histological examinations also revealed degenerative changes and inflammatory foci in the colon of DEN-treated rats. These observations are in line with our findings that DEN caused a significant decrease in crypt depth and a trend towards increased PCNA staining and Alcian blue staining. More research is needed to delineate whether these effects were induced by the DEN-injections or were a result of the hepatocarcinogenesis, and to what extent chemotherapy aggravates these.

In our study, we found the limited efficacy of DOX and IDA in reducing tumor burden in a DEN-induced HCC murine *in vivo* model. It is important to note that timing and dosing may have played a role in interpreting the effects of anthracyclines in our DEN-induced HCC mouse model. Due to the weight loss observed in the mice receiving IDA or DOX, we limited the treatment to three weeks, capturing only the acute phase of the drug’s impact. It is possible that prolonged, repeated or increased exposure to these agents would have increased the anti-tumor effect, amplified fibrosis and/or induced more damage to the intestinal mucosa. While IDA exhibited higher cytotoxicity *in vitro*, its promotion of HSC activation and pro-inflammatory signaling *in vivo* raises concerns and underscores the need for further research into the temporal and microenvironmental effects of anthracyclines. Furthermore, the study highlights the importance of considering chemotherapy-induced side-effects, as both anthracyclines contributed to weight loss and possibly induced damage to the intestinal crypts, which would affect overall treatment tolerance and wellbeing. These findings underscore the complexity of anthracycline-based chemotherapy in HCC and emphasize the need for further research into refining therapeutic strategies that balance drug response with minimizing adverse effects. Future studies should explore alternative chemotherapeutic agents, combination therapies, and the temporal dynamics of drug-induced inflammation and fibrosis to improve treatment outcomes for patients with unresectable intermediate-stage HCC, while also taking in account potential side-effects.

---

**Supplementary Material Table 1.**
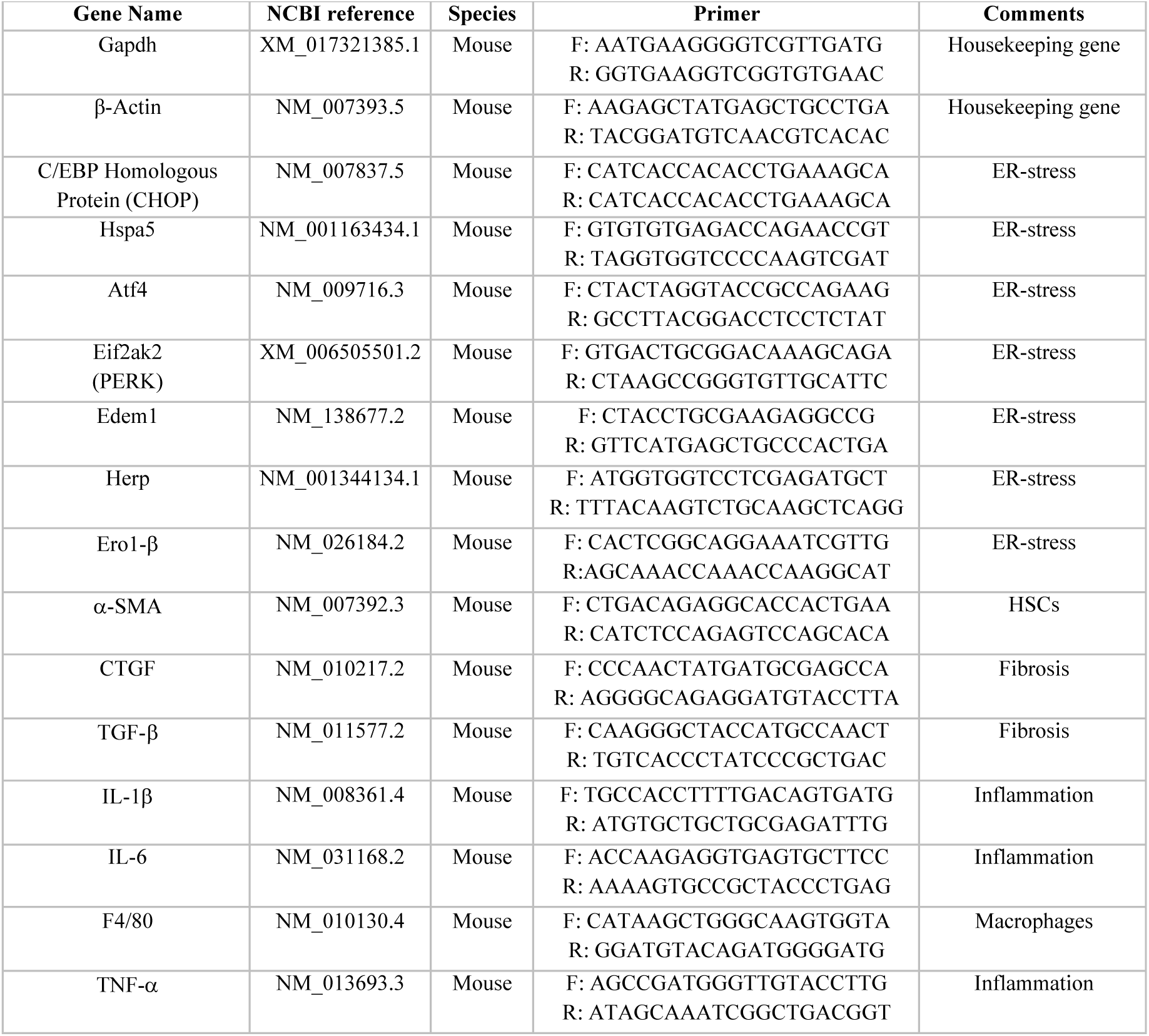

## Notes

### Competing Interest Statement

The authors have declared no competing interest.

